# Unbiased MD simulations characterize lipid binding to lipid transfer proteins

**DOI:** 10.1101/2023.11.24.568550

**Authors:** Sriraksha Srinivasan, Daniel Alvarez Lorenzo, Stefano Vanni

## Abstract

The molecular characterization of lipid binding to lipid transfer proteins (LTPs) is fundamental to the understanding of several aspects of their mechanistic mode of action. However, obtaining lipid-bound structures of LTPs is not straightforward owing to caveats in current experimental structural biology approaches. As a result, several structures of LTPs, and most notably almost all of those that have been proposed to act as bridges between membrane organelles, do not provide the precise location of their endogenous lipid ligands. To address this limitation, computational approaches are a powerful alternative methodology, but they are often limited by the high flexibility of lipid substrates. In this work, we develop an *in silico* protocol based on unbiased coarse grain molecular simulations in which lipids placed in bulk solvent away from the protein can spontaneously bind to LTPs. This approach accurately determines binding pockets in LTPs and provides a working hypothesis for the pathway via which lipids enter LTPs. We apply this approach to characterize lipid binding to bridge-like LTPs belonging to the Vps13-Atg2 family, for which the lipid localization inside the protein is currently unknown. Overall, our work paves the way to determine binding pockets and entry pathways for several LTPs in an inexpensive, fast, and accurate manner.

## Introduction

Eukaryotic cells are organized into separate membrane-bound organelles, each with a characteristic lipid composition necessary for its proper functioning^1^. To achieve this complex homeostatic balance, lipids need to be selectively mobilized between different organelles in response to both extra-and intra-cellular stimuli^2^.

Intracellular lipid trafficking is largely achieved by two mechanisms – vesicular and non-vesicular transport. Non-vesicular lipid trafficking is mediated by a large group of lipid transfer proteins (LTPs), which encapsulate lipids in the hydrophobic cavity of a soluble lipid transfer domain (LTD) and transfer them between membranes of different organelles within the cell^3, 4^. Notably, many LTPs have been discovered in the last few years^5–7^, suggesting that intracellular lipid transport is more widespread and prominent than originally thought. Concomitantly with the identification of new LTPs, new mechanisms of lipid transport are also being proposed^4, 8^. Notably, certain LTPs have been suggested to transfer lipids between organelles by physically bridging them at membrane contact sites (MCS) and by establishing long hydrophobic tunnels between the two organellar membranes^3, 4, 9,10^.

Despite this plethora of new data on LTP cellular functions^3, 4, 11^, a detailed understanding of their mechanistic mode of lipid transport remains limited. For example, even when a high-resolution structure of the LTP is available, the co-transported lipids are often missing, possibly as a consequence of the dynamic behavior of lipids inside the protein cavity. In addition, only few high-resolution structures of LTPs are available, and especially for those proposed to work via a bridge-like mechanism, AlphaFold-derived models are often used to provide a mechanistic interpretation of the functional data^12–22^. Finally, basic mechanistic features such as the exact entry/exit pathways for the loading and unloading of the lipids from the protein remain largely unknown.

To overcome these limitations, computational approaches hold promise to identify lipid binding poses for LTPs. A classical strategy in this regard is molecular docking^23^. While this technique is widely used to determine ligand binding pockets in proteins^24^, the high flexibility of lipid chains, possibly also inside the protein binding cavity, results in a major challenge in the case of LTPs. Recently, promising attempts to use AI-derived tools to obtain the structure of protein-ligand complexes by directly building the protein structures around the corresponding molecules have been proposed^25^, but these methods still perform quite poorly for low-affinity ligands^25^. Alternatively, both unbiased atomistic molecular dynamics (MD) simulations, in which the ligand of interest is placed in the bulk solvent and eventually binds to the protein^26^, as well as enhanced sampling MD methods^27 28^, have been used to determine binding pathways for small molecules^29, 30^. However, since these processes occur over relatively long timescales, these approaches are inherently computationally expensive and thus difficult to extend to high-throughput investigations. To alleviate these computational bottlenecks, a promising computational protocol based on unbiased coarse-grain (CG) MD simulations to determine protein-ligand interactions for small drug-like molecules has been recently proposed^31^. There, the authors elegantly demonstrate that chemical-specific CG simulations (using the MARTINI force field^32^) can be used to simulate protein-ligand binding using a brute force approach, paving the way to potentially screening numerous drugs and proteins in a high-throughput fashion^31^.

In this work, we explore the possibility of extending this protocol to the complex cases of lipids in LTPs. By tuning the computational parameters, we succeed in observing the spontaneous binding of lipids to LTPs and their eventual entry into the experimentally determined lipid binding pocket whenever this structural information is available. Our approach identifies a dominant entry pathway of lipids into LTPs and provides a structural view of the lipid density inside bridge-like LTPs (BLTPs), such as Vps13 and Atg2, that have been proposed to act as bulk lipid transporters but for which the lipid localization inside the cavity has remained uncharacterized so far. Our results pave the way for an improved characterization of the molecular mode of action of LTPs, and potentially for the *in silico* identification of new ones, in a high-throughput fashion at minimal computational cost.

## Results

### Brute force unbiased coarse-grain molecular dynamics simulations can reproduce crystallographic poses of lipids bound to LTPs

To determine whether unbiased CG-MD simulations can correctly identify lipid-protein interactions for LTPs, we first adapted a protein-ligand protocol recently proposed for small drug-like molecules using the CG MARTINI force field^31^. In this protocol, drug-like molecules are placed in bulk solvent in the presence of a target protein, and during unbiased MD simulations they can spontaneously relocate into the protein binding pocket, due to the smoothness of the free energy profile in the CG representation^33^. We reasoned that a similar protocol could work for lipid binding to LTPs even if lipid molecules carry extremely hydrophobic moieties: in the absence of alternative hydrophobic binding partners (such as a lipid bilayer) the lipid of interest could potentially be able to find the protein cavity of LTPs within reasonable time scales.

To test this hypothesis, we first identified several proteins that have been co-crystallized with a single lipid they are proposed to transport (Table 1, rows 1 to 10). These include Ceramide-1-phosphate transfer protein (CPTP), Fatty acid binding protein-1 (FABP1), Mitoguardin-2 (Miga2), Maintenance of lipid asymmetry protein C (MlaC), Phosphatidyl-choline transfer protein (PCTP), Neurofibromin-1 (NF1), Oxysterol binding protein homolog-6 (Osh6), Phosphatidyl-inositol transfer protein-alpha (PITP-A), Chromatin structure-remodeling complex subunit Sfh1, and START domain-containing protein-11 (Stard11, also called ceramide transfer protein (CERT)). Starting from the X-ray structures, we next set up the following protocol: each protein was stripped off its co-crystallized ligand(s), placed in a box of water with 0.12M NaCl with a lipid randomly inserted into the bulk solvent, and 1 μs-long unbiased MD simulations were performed (Fig. 1A and Movie S1). For each protein, 5 independent replicas were carried out. Hydrophobic tails of all simulated lipids were modeled as oleoyl tails (18:1), while head groups were chosen to be similar to the ones in the corresponding crystal structure of the protein. In case parameters for a given lipid in the crystal structure were unavailable in the MARTINI force field, structurally similar head groups were chosen for comparison. For example, 1,2-dioleoyl-sn-glycero-3-phosphate (DOPA) was used instead of ceramide-1-phosphate.

**Figure 1:**
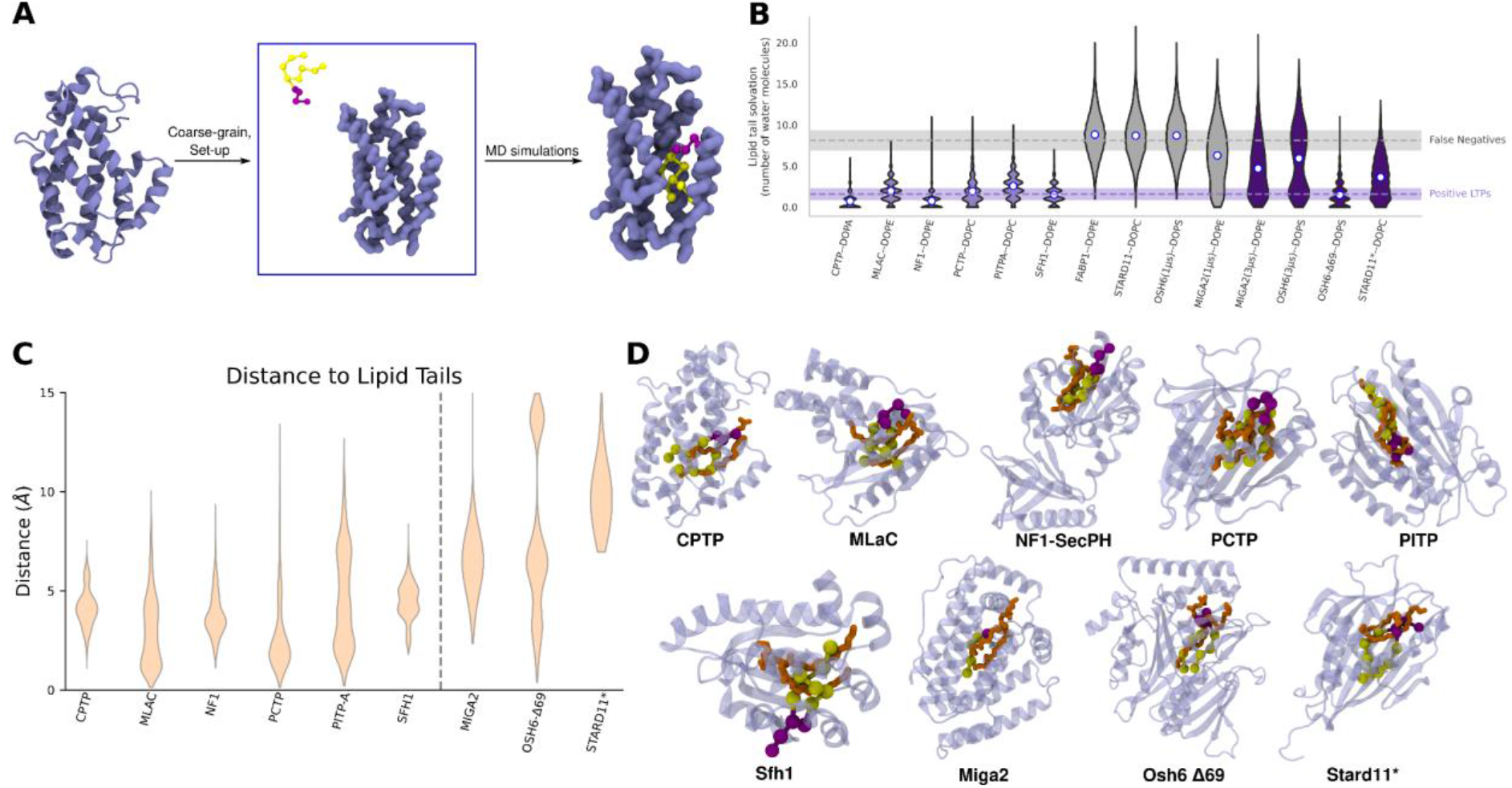
CG simulations reproduce the correct lipid binding pose to LTPs. **(A)** Unbiased CG-MD simulation protocol used in this study. Atomistic structures of the protein were coarse-grained and set-up with a lipid randomly placed in a bulk solvent far away from the protein, followed by MD simulations. **(B)** Lipid tail solvation in the last 100 ns of the CG-MD trajectories. Light purple indicates true positives, grey indicates false negatives, and dark purple indicates true positives after modification of our initial protocol. Dots at the center of the violin plot indicate average values for each protein while the two dotted lines indicate the average lipid tail solvation for the true positives (purple, 6/10 proteins) and the false negatives (gray, 4/10 proteins). **(C)** Distance between the center of mass of the hydrophobic tails of the experimental ligands and the lipids tested in the simulations. Only the frames with a lipid tail solvation lower than 2 were considered. **(D)** Illustrative examples of agreement between lipid positions arising from our protocol (lipid tails: yellow, headgroups: purple) and the lipid/ligand position in X-ray structures (orange). The use of a different force constant for the elastic network of Stard11 is indicated by a *.

**Table 1.**
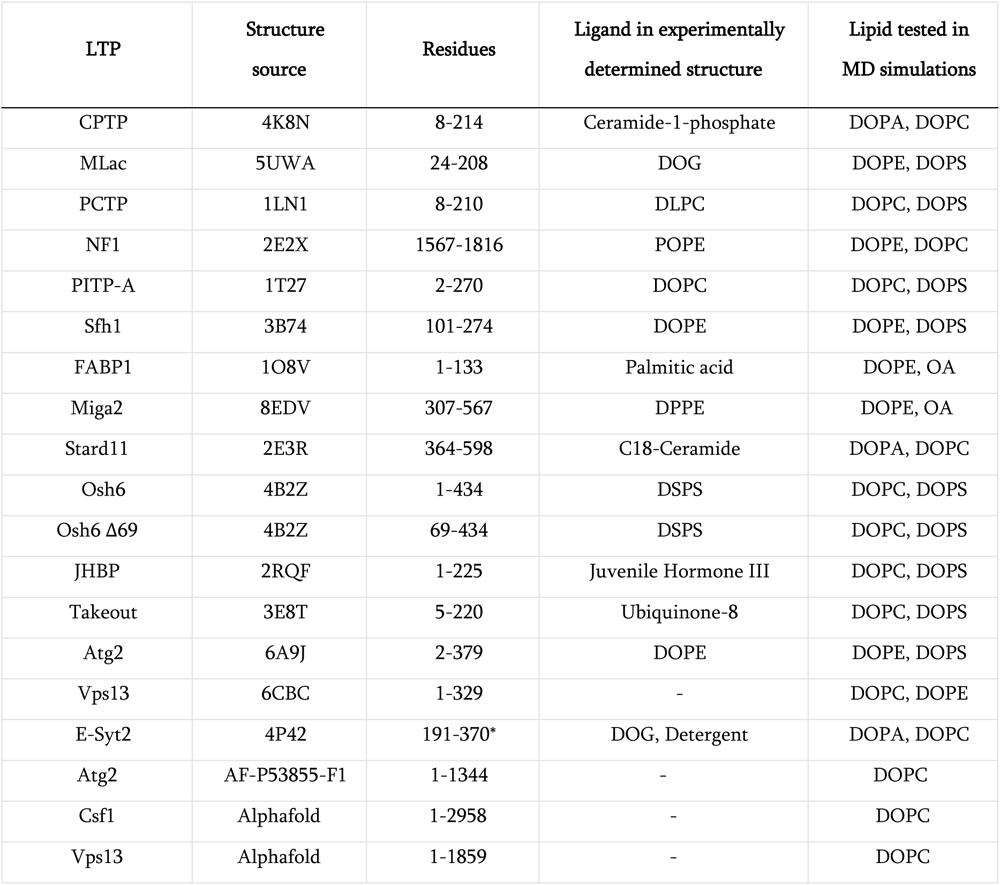
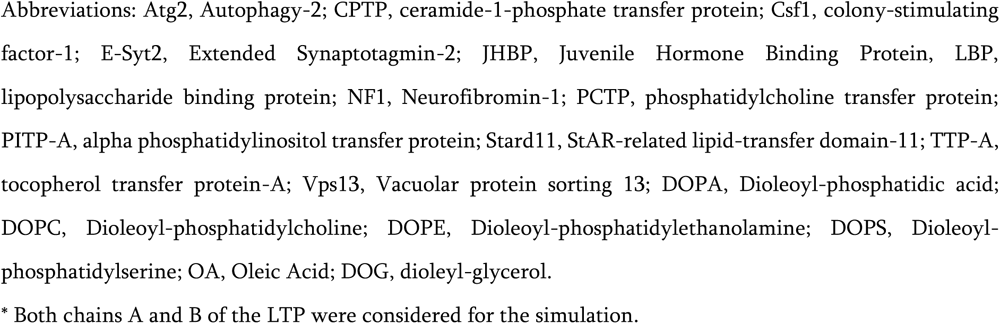
Details of the LTPs tested in this study.

| LTP | Structure source | Residues | Ligand in experimentally determined structure | Lipid tested in MD simulations |
| --- | --- | --- | --- | --- |
| CPTP | 4K8N | 8-214 | Ceramide-1-phosphate | DOPA, DOPC |
| MLac | 5UWA | 24-208 | DOG | DOPE, DOPS |
| PCTP | 1LN1 | 8-210 | DLPC | DOPC, DOPS |
| NF1 | 2E2X | 1567-1816 | POPE | DOPE, DOPC |
| PITP-A | 1T27 | 2-270 | DOPC | DOPC, DOPS |
| Sfh1 | 3B74 | 101-274 | DOPE | DOPE, DOPS |
| FABP1 | 1O8V | 1-133 | Palmitic acid | DOPE, OA |
| Miga2 | 8EDV | 307-567 | DPPE | DOPE, OA |
| Stard11 | 2E3R | 364-598 | C18-Ceramide | DOPA, DOPC |
| Osh6 | 4B2Z | 1-434 | DSPS | DOPC, DOPS |
| Osh6 Δ69 | 4B2Z | 69-434 | DSPS | DOPC, DOPS |
| JHBP | 2RQF | 1-225 | Juvenile Hormone III | DOPC, DOPS |
| Takeout | 3E8T | 5-220 | Ubiquinone-8 | DOPC, DOPS |
| Atg2 | 6A9J | 2-379 | DOPE | DOPE, DOPS |
| Vps13 | 6CBC | 1-329 | - | DOPC, DOPE |
| E-Syt2 | 4P42 | 191-370* | DOG, Detergent | DOPA, DOPC |
| Atg2 | AF-P53855-F1 | 1-1344 | - | DOPC |
| Csf1 | AlphaFold | 1-2958 | - | DOPC |
| Vps13 | AlphaFold | 1-1859 | - | DOPC |

Abbreviations: Atg2, Autophagy-2; CPTP, ceramide-1-phosphate transfer protein; Csf1, colony-stimulating factor-1; E-Syt2, Extended Synaptotagmin-2; JHBP, Juvenile Hormone Binding Protein, LBP, lipopolysaccharide binding protein; NF1, Neurofibromin-1; PCTP, phosphatidylcholine transfer protein; PITP-A, alpha phosphatidylinositol transfer protein; Stard11, StAR-related lipid-transfer domain-11; TTP-A, tocopherol transfer protein-A; Vps13, Vacuolar protein sorting 13; DOPA, Dioleoyl-phosphatidic acid; DOPC, Dioleoyl-phosphatidylcholine; DOPE, Dioleoyl-phosphatidylethanolamine; DOPS, Dioleoyl-phosphatidylserine; OA, Oleic Acid; DOG, dioleoyl-glycerol.
\* Both chains A and B of the LTP were considered for the simulation.

To assess lipid binding, we used two distinct metrics. First, we collected the time traces of the minimum distance between the protein and the lipid (Fig. S1). Second, to discriminate between *bona fide* binding events inside the lipid binding pocket versus futile lipid-binding events on the external surface of the protein, we used as a metric the solvation number of the lipid tails (defined as the number of water molecules within 5 Å of the lipid acyl chains) in the last 100 ns of the trajectory of each replica (Fig. 1B). We selected this property since a lipid bound to an external surface of the protein would be surrounded by a larger number of water molecules as opposed to lipids bound to a hydrophobic cavity.

For the 10 proteins we tested, we observed two distinct behaviors: in some cases (6/10: CPTP, MLac, NF1, PCTP, PITP-A, Sfh1; Fig. 1B) the lipid could always bind spontaneously to the protein of interest, often irreversibly, as indicated by the time traces of the minimum distances between the protein and the lipid (Fig. S1). In those cases, solvation analysis indicates that the lipid tails are desolvated at the end of the trajectory, thus residing inside a hydrophobic binding pocket (Fig. 1B, light purple).

In other cases (3/10: FABP1, Stard11, Osh6), no entry of the lipid was observed (Fig. S1 and 1B, gray traces), and the lipid could not find the binding pocket within the 1 μs elapsed simulation time. In one case (Miga2) the lipid entered the binding pocket only in 3 of the 5 replicas (Fig. S1 and Fig. 1B).

Next, we focused our attention on the false negative results (Fig. 1B). We reasoned that the origin of such behaviour could be related to high free energy barriers originating from protein dynamics or conformational changes potentially associated to lipid entry in physiological conditions and that are not well-reproduced by our CG simulations. To qualitatively test this hypothesis, we perform three tests. First, we extended our simulations from 1 to 3 μs. This approach improved our results for Osh6 (Fig. S1), with two replicas displaying lipid binding in the extended simulation, and for Miga2, in which we observed lipid binding in one additional replica (Fig. S1 and 1B). Next, we tested our protocol on lid-less Osh6 (Osh6 Δ69), since oxysterol related proteins (ORP) domains, like other LTDs, are known to have a lid-like helical region at the N-terminus that regulates lipid entry into the protein^34, 35^. Indeed, while we observed lipid entry for full length Osh6 in two replicas (in 3 μs), all five replicas displayed lipid entry for Osh6 Δ69 (Fig. 1B, S1). Third, we opted to decrease the elastic network force constant that keeps the protein conformation close to its crystallographic structure in CG simulations, thus promoting increased flexibility. Indeed, upon reduction of the force constant (from 1000 to 300 kJ mol^-1^ nm^-2^) we could observe spontaneous lipid entry in the cavity of Stard11 (Fig. 1B, S1), but not for FABP1 (Fig. S1).

Finally, to test whether the de-solvated lipid-binding pose we observed is consistent with the one determined by X-ray crystallography, for the 9 proteins out of 10 where we could observe spontaneous binding (Fig. 1B), including after modification of the initial protocol, we computed the distance between the center of mass of the hydrophobic tails in the experimental structure and the corresponding one of the CG lipids in the last 100 ns of the simulation trajectories, exclusively for MD frames with a low lipid tail solvation (<2 water molecules, consistent with the mean value over the positive LTPs of 1.8 molecules on average) (Fig. 1C).

Even though our simulations display a large variability, in all cases we could observe values very close to the X-ray distance (Fig. 1C, D), thus identifying binding poses very close to the crystallographic structure, with the sole exception of STARD11, where CG-MD simulations suggest a lipid binding pose with the lipid tails located more deeply inside the protein cavity than in the corresponding structure from X-ray crystallography (Fig. 1D). The observation that our protocol provides additional de-solvated lipid poses with respect to the crystal structure is potentially interesting. On the one hand, this observation could be simply attributed to the inaccuracy of our CG approach. On the other hand, this observation could originate from the ability of our protocol to identify additional lipid binding poses. These conformations might be important along the lipid entry/exit pathway and their presence is consistent with the hypothesis that the lipid must not be trapped in highly stable states inside the pocket in order to facilitate its uptake and release from membranes by the LTPs.

### CG-MD simulations cannot reproduce the experimentally-determined lipid specificity of LTPs

Next, we investigated whether our assay could reproduce the experimentally determined sensitivity for specific lipids. To do so, we performed identical simulations with a second lipid that is not known to bind to the tested LTPs. In all cases, we found an identical binding behaviour (Fig. S1), indicating that our protocol is unable to discriminate between lipids with different headgroups. This is also true when lipids with significantly different sterical hindrances were tested, as is the case for oleic acid in Miga2 and FABP1 (for both proteins fatty acids have been proposed to be a natural substrate)^36, 37^. Finally, analysis of the polar heads placement in our simulations reveals a large variability in the binding poses between replicas (Fig. S2), as a consequence of the lack of sensitivity for the different head groups. Rather, the polar heads can be stabilized by both interaction with the protein or by the surrounding solvent in our CG simulations.

Furthermore, to investigate the ability of our protocol to discriminate between different lipids, we applied it to 2 proteins, the takeout protein and the juvenile hormone binding protein (JHBP), that possess Synaptotagmin-like mitochondrial-lipid-binding (SMP) domains that are structurally similar to that of known LTPs. The takeout protein and the JHBP do not transport lipids, but they transport hydrophobic, lipid-like molecules: ubiquinone-8 and juvenile hormone III respectively. In the presence of 1,2-dioleoyl-sn-glycero-3-phosphocholine (DOPC) and 1,2-dioleoyl-sn-glycero-3-phospho-L-serine (DOPS) lipids, we observed spontaneous binding and very modest lipid solvation for both proteins (Fig. S3). Taken together, these results indicate that while our approach succeeds in the identification of hydrophobic lipid-binding pockets for LTPs, it is not able to recapitulate the experimental selectivity for specific lipids, and thus cannot be used to interrogate lipid specificity of LTPs.

### Unbiased CG-MD simulations can discriminate between bona-fide LTPs and negative control proteins that contain non lipid-specific large cavities

The observation that our protocol is sometimes unable to reproduce the experimentally determined binding of lipids to LTPs suggests that our approach might not be biased towards false positive results. To further stress-test our protocol in this direction, we next investigated whether we could correctly flag proteins that do not transport lipids despite the presence of a large cavity in their structure. To do so, we selected 10 proteins that are not known to bind lipids but possess a cavity of large volume^38^ (Table 2), that is either hydrophobic or hydrophilic, as negative controls. Lipid binding to these negative control proteins was assessed using the same protocol describe above, using DOPC as a model lipid.

**Table 2.** Details of the negative control proteins tested in this study.

| Negative control | PDB ID | Nature of cavity | Lipid tested in MD simulations |
| --- | --- | --- | --- |
| Ferric-citrate transporter | 1PO0 | Hydrophilic | DOPC |
| Sheep lactoperoxidase | 2IKC | Hydrophilic | DOPC |
| Hydroxylase-Regulatory Protein Complex | 2INP | Hydrophilic | DOPC |
| Sensory Rhodopsin II | 1JGJ | Hydrophobic | DOPC |
| Na<sup>+</sup>/H<sup>+</sup> antiporter NhaA | 1ZCD | Hydrophobic | DOPC |
| T4-Lysozyme | 181L | Neutral | DOPC |
| CDK2 | 1AQ1 | Hydrophobic | DOPC |
| Concanavalin A | 3ENR | Hydrophilic | DOPC |
| Thyroid Hormone Receptor | 1NAX | Hydrophobic | DOPC |
| Transcription Factor T | 1XBR | Hydrophilic | DOPC |

For these negative controls, the lipids bound stably to the proteins during the CG simulations in many instances (Fig. S4). However, upon computing the spatial density of the lipids during the last 100 ns of the simulation trajectory, a more outspread occupancy map was observed for the negative control proteins (Fig. 2A). This observation suggests that non-specific interactions of the lipid with different exterior surfaces of the proteins, rather than specific interactions with a lipid binding pocket, characterize lipid binding with the negative controls in our assay. To quantify this observation, calculation of the solvation number of the lipid indicates that lipid solvation was significantly higher in the case of the negative controls (mean value = 6.4) than that in the positive-result LTPs (mean value = 1.8, Fig. 2B). This indicates that, for negative controls, lipids bound almost exclusively to external surfaces of the proteins despite the presence of a cavity. Precisely, out of 50 control simulations (5 for each protein), we observed a single exception in one of the replicas of concanavalin A (PDB ID: 3ENR, Fig. 2B), further suggesting that our approach is quite robust against false positive results.

**Figure 2:**
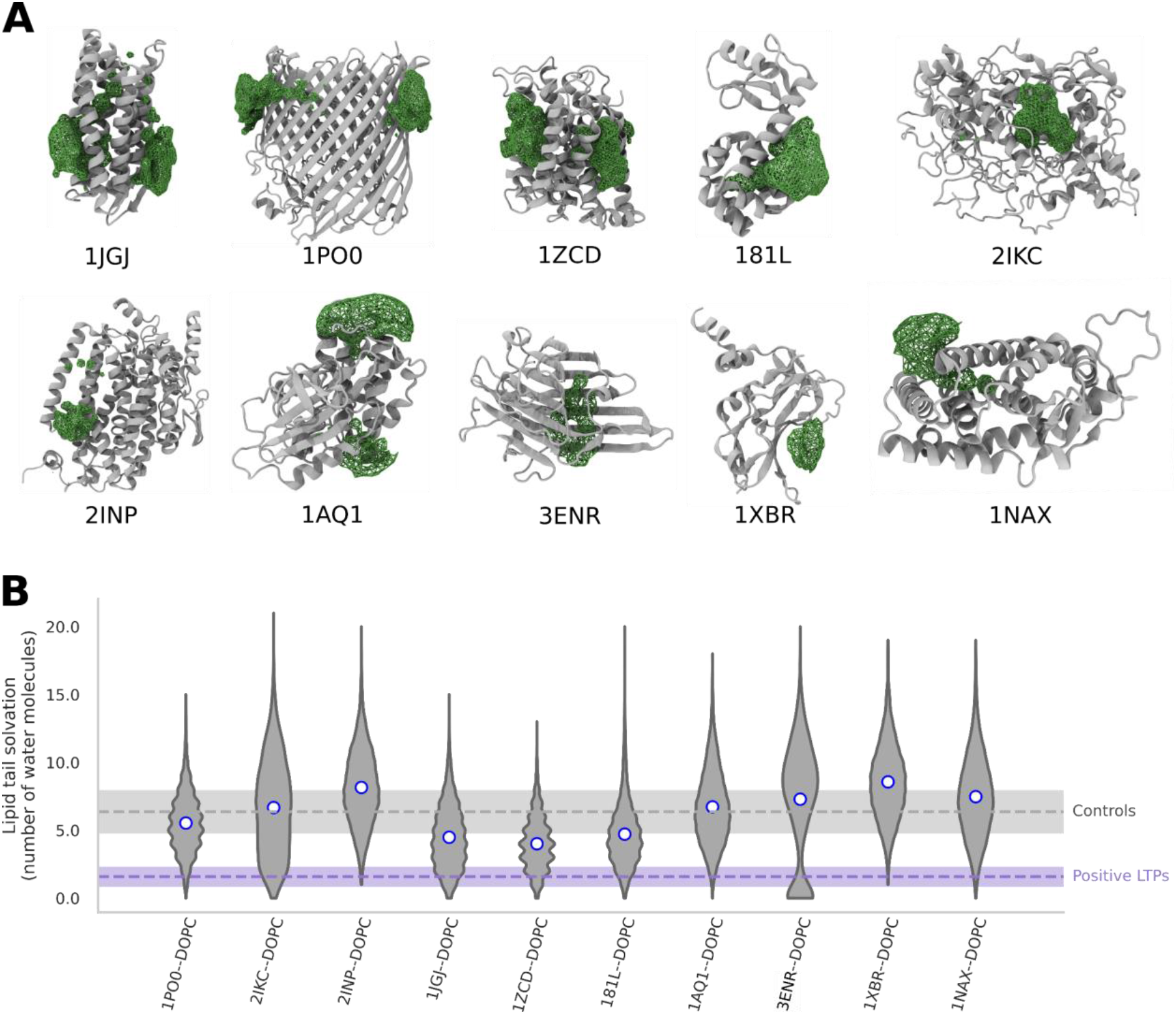
CG simulations do not show lipid binding to non-LTPs. **(A)** Spatial density maps of lipid occupancy in the last 100 ns of unbiased CG simulations with DOPC for non-LTP negative controls. **(B)** Lipid solvation of non-LTPs in the last 100 ns of the trajectory. Dots indicate average values.

### Entry of the lipid into the protein cavity occurs via a dominant pathway

We next focused our attention on the path of lipid entry inside the LTP cavity, as this process is critical to understanding their mechanism of action. Since our protocol does not describe the physiological conditions of lipid entry/exit, with the lipid initially placed in bulk solvent rather than in a lipid bilayer, we wanted to investigate whether lipid entry was following a single dominant, possibly physiological, pathway or if rather the protein structure provided several entry points for fully water-solvated lipids.

Visualization of the simulation trajectories revealed the presence of a dominant pathway of lipid entry for all LTPs. By measuring the fraction of replicas in which the corresponding pathway was observed, we noticed that this pathway was unique for some LTPs, such as CPTP, PITPA, and Osh6 Δ69 (Fig. 3A), and observed with a probability between 67 and 90% for all other LTPs. The pathway observed in these simulations also corresponds to the one that has been proposed in literature for some proteins such as the CPTP^39^, Osh6^35^ and members of the StardD family^40^, to which PCTP and Stard11 belong.

**Figure 3:**
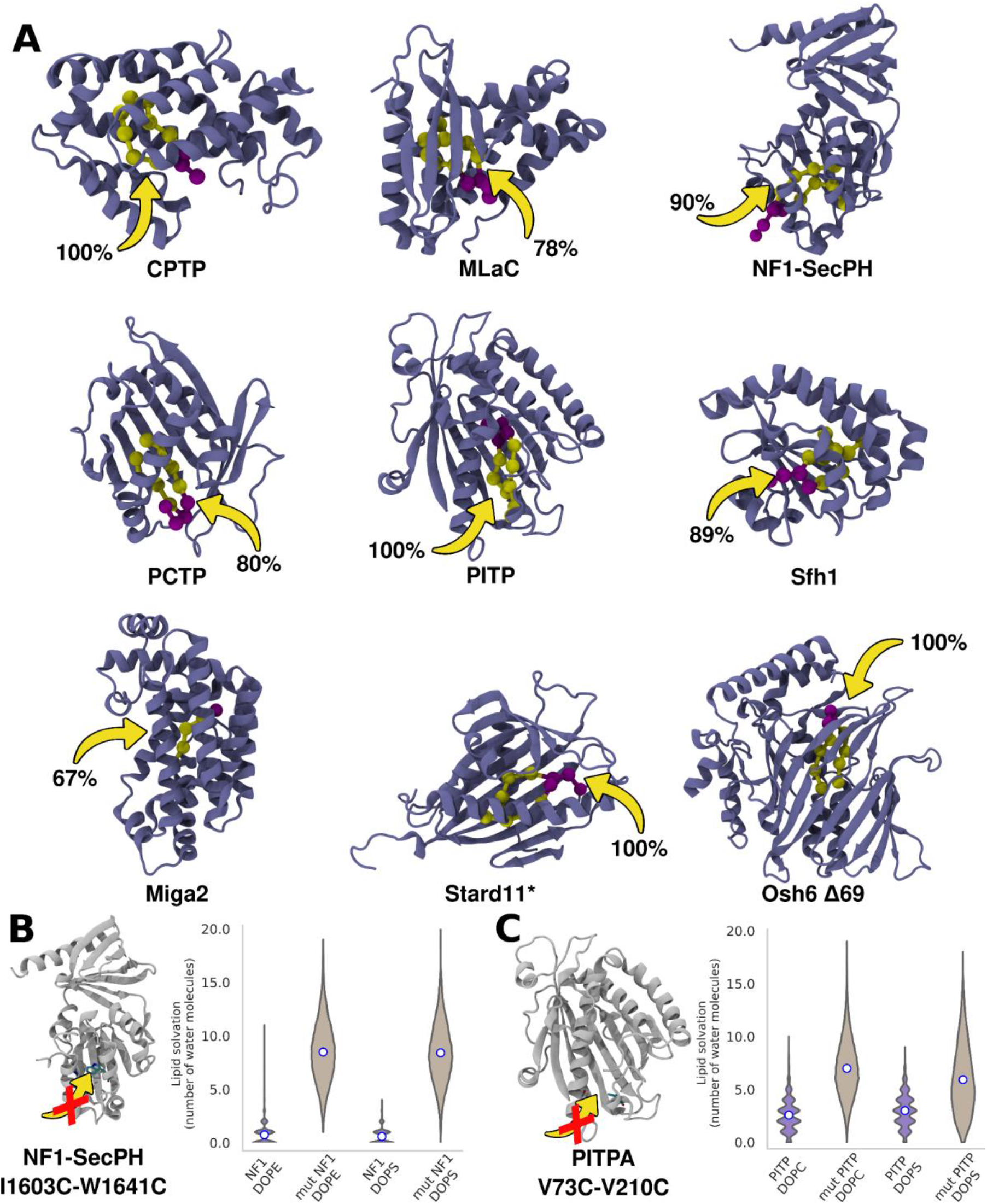
Lipid entry pathways: **(A)** Entry pathway of lipids into the LTD. Percentage value beside arrows indicate the fraction of replicas in which the pathway was observed. Protein represented in cartoon representation and the simulated lipid with tails in yellow and the headgroup in purple. **(B,C)** Disulfide bridges along the lipid-entry path in NF1-Sec PH and PITP-A abolish lipid entry into the binding pocket as indicated by the lipid solvation plots. The use of a different force constant for the elastic network of Stard11 is indicated by a *.

Finally, to test the robustness of the entry pathway we observed, we designed mutations to block this specific entry process. To do so, we performed identical simulations after introducing disulfide bridge mutations along the lipid entry pathway - I1603C-W1641C in the case of NF1-SecPH, and V73C-210C in the case of PITP (Fig. 3B,C). The residues were chosen such that the distance between Cα atoms of the residues is less than 8Å. We observed that in both cases, the presence of the disulfide bridge abolished lipid entry into the protein entirely, as shown by the lipid solvation in Fig. 3B,C.

### Brute-force CG-MD simulations can characterize the lipid-binding cavity of poorly characterized lipid transport proteins

As potential applications of our protocol, we first focused on proteins that have been proposed to transport lipids, but for which no experimental lipid-bound structure was determined. These include, for example, the recently characterized yeast ceramide transporter Svf1^41^, SNX25^13^, which has been proposed to belong to a new class of LTPs, and the repeating beta groove (RBG) proteins Mdm31 and AsmA^19^. Using the AF structure as a starting point, our protocol is indeed able to propose a lipid binding pose for these proteins (Fig. 4A-E).

**Figure 4:**
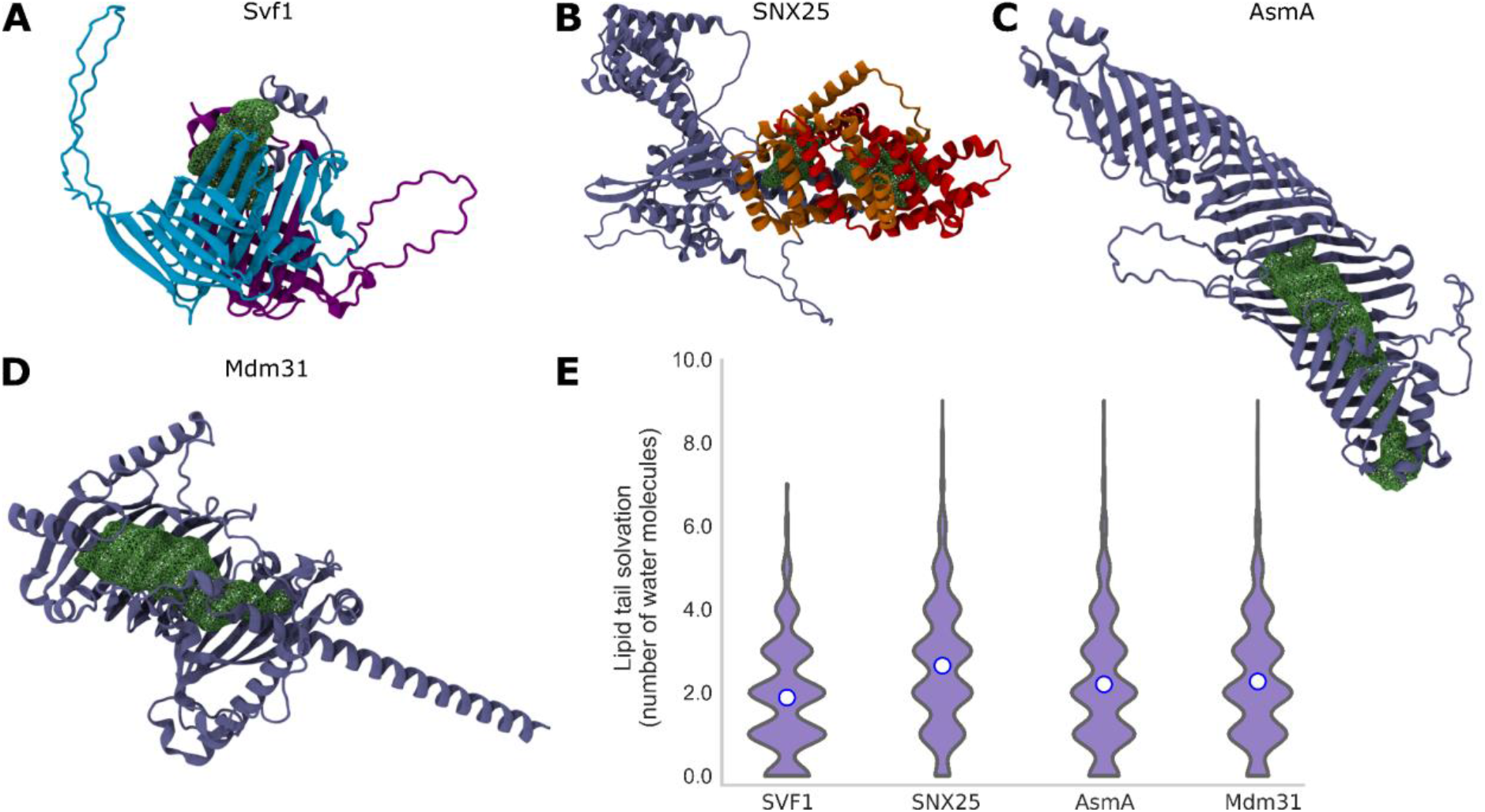
Identification of the hydrophobic cavity of potential LTPs. Spatial density maps **(A-D)** of: **(A)** Svf1, **(B)** SNX25 (B), **(C)** AsmA, and **(D)** Mdm31. The two lipocalin domains of Svf1 are shown in blue and violet. The PXA and PXC domains of SNX25 are displayed in red and orange, respectively. Averaged lipid densities are shown in green. **(E)** Lipid tail solvation for the four potential LTPs, only the replicas with lipid binding were considered for the analysis.

The identified lipid binding pocket of Svf1 is located between the two lipocalin domains (blue and violet in Fig. 4A), which is in good agreement with what has been recently suggested using blind docking^41^. For SNX25, our protocol suggests that lipid binding, which occurs in 4 out of 5 replicas, takes place in a long and conserved hydrophobic pocket formed by the PXA and PXC domains (red and orange in Fig. 4B, respectively), in agreement with what was recently proposed based on structural considerations^13^. AsmA and Mdm31 are two prokaryotic RBG proteins distantly related to the eukaryotic bridge-like LTP (BLTP) superfamily, which includes well studied LTPs such as Vps13 and Atg2. As expected, our protocol displays lipid binding in all replicas (AsmA) and in 3 out of 5 replicas (Mdm31), within the hydrophobic cavity formed by the RBG domains in a similar manner as their eukaryotic counterparts (Fig. 4C,D).

In summary, our protocol can identify potential hydrophobic pockets of LTPs that are poorly characterized at the structural level, and our results are in good agreement with alternative methods such as docking or structural analysis.

### LTPs that can bind multiple lipids in their cavity possess several continuous lipid interacting regions

Next, we applied our protocol to BLTPs^19, 42^. Recently, many LTPs have been proposed to transport lipids via a bridge-like mechanism by establishing a continuous hydrophobic conduit between membrane organelles^4, 19^. Within this model, LTPs could bind many lipids concomitantly, and they could contribute to bulk lipid transport between organelles, such as in autophagosome formation^43, 44^ or lysosomal repair^45, 46^. Yet, available high-or medium-resolution structures of BLTPs are scarce and entirely devoid of lipids, with the sole exception of one single phosphatidylethanolamine (PE) molecule in the structure of a small region of ATG2^43^.

To investigate lipid occupancy in BLTPs, we adapted our protocol to work in the presence of multiple lipids. To do so, we performed lipid-addition simulations iteratively, to avoid lipid-lipid interactions (leading to micelle formation) in the solvent before binding to the protein. As a first test to validate our protocol, we investigated the Synaptotagmin-like Mitochondrial-lipid-binding Protein (SMP) domain of E-Syt2 dimer^47–49^. While this protein is proposed to work via a shuttle-like mechanism^47, 50^, it has been co-crystallized with multiple lipids in its cavity (2 diacylglycerols and 2 detergents), providing a natural positive control for our protocol.

Indeed, we observed that with the iterative lipid addition process, at least 5 lipids can be accommodated in the cavity of the SMP domain as indicated by the low solvation of the lipid tails (Fig. 5D). Intriguingly, the average spatial density of lipids in E-Syt2 spans the entire hydrophobic conduit rather than a specific lipid binding pocket (Fig. 5A).

**Figure 5:**
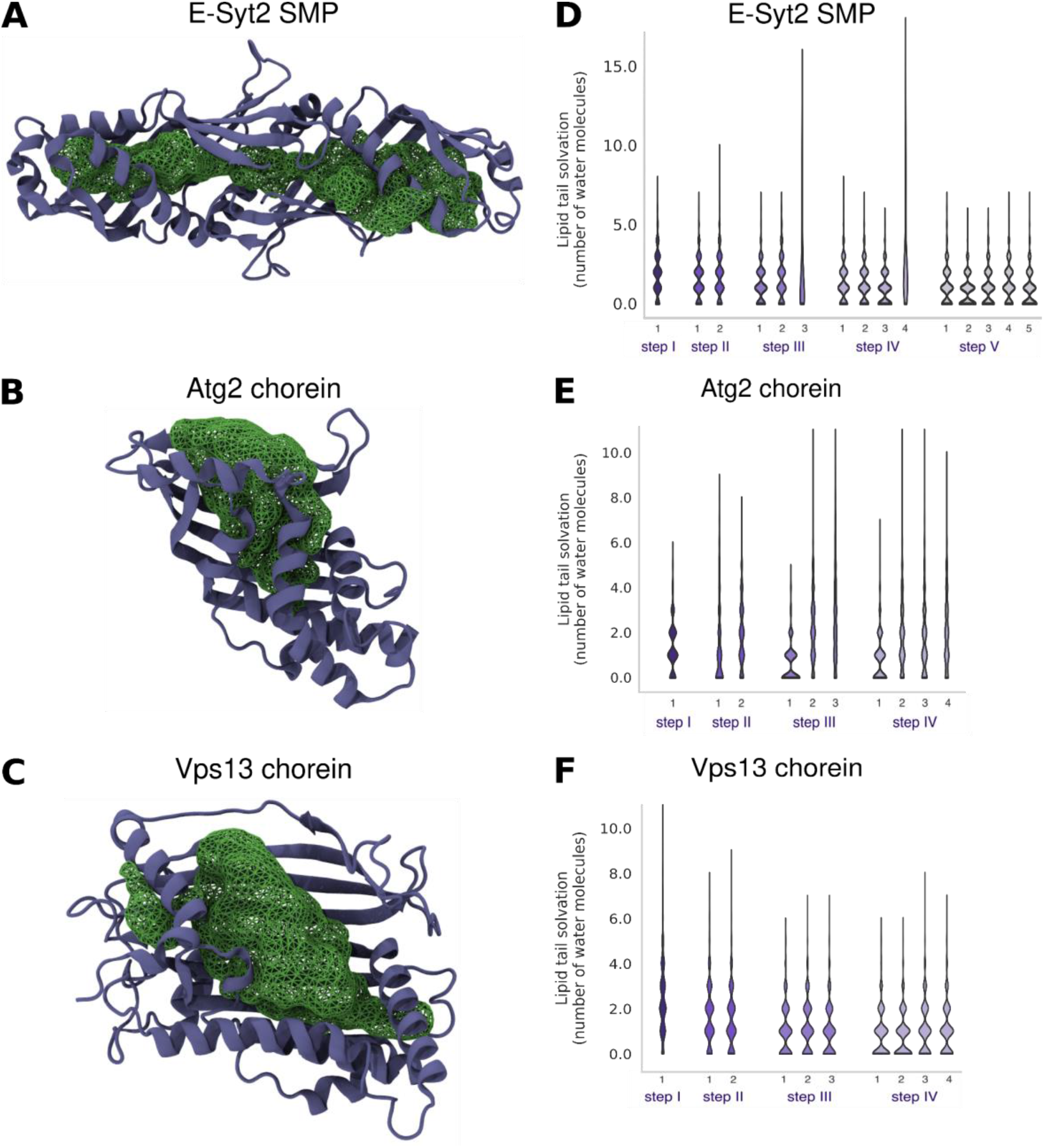
LTDs with multiple lipid binding regions. (**A-C**) Spatial density maps of lipids and (**D-F**) corresponding solvation for each lipid in the multi-step iterative protocol for the **(A, D)** SMP domain of E-Syt2, **(B, E)** Chorein motif of Atg2, and **(C, F)** Chorein motif of Vps13. The protein is shown in purple; the averaged lipid density in green.

We next applied our protocol to Atg2 and Vps13, as these proteins have been proposed to bind multiple lipids at once. We initially restricted our analysis to the respective chorein domains of these proteins, as these regions have been solved at high resolution (2.7 and 3.0Å, respectively^43, 51^). Again, our protocol shows that the proteins are able to bind multiple lipids and form a continuous lipid-filled tunnel spanning the entirety of the hydrophobic cavity formed by the repeating beta groove (RBG) domains (Fig. 5B,C).

Taken together, these data indicate that our protocol is able to predict the simultaneous binding of multiple lipids to BLTPs, and it confirms that the lipid binding mode for these proteins is distinct from those proposed to work in a shuttle-like fashion. Of note, in all cases we observed that the presence of multiple lipids further decreases their solvation (Fig. 5D-F). This suggests that the presence of lipids inside the binding cavity could promote sequestration of lipids from the lipid bilayer and, hence, lipid transport.

Next, we investigated multiple proteins that belong to the BLTP superfamily, Csf1 (also known as BLTP1 or Tweek), Vps13 and Atg2 in their entirety (Fig. 6A-C). For Atg2, we used the Alphafold structure (identifier AF-P53855-F1) and residues 1 to 1344, as the C-terminal region consists of several helices with low pLDDT score that are not part of the hydrophobic cavity. In the case of Csf1, the structure of three different fragments was predicted using Alphafold and then aligned to get the whole protein structure (2958 residues), which was considered entirely. Finally, for Vps13, a similar protocol was employed, but residues 1860-3144 were discarded, as they do not belong to the extended-Chorein domain that forms the hydrophobic cavity.

**Figure 6:**
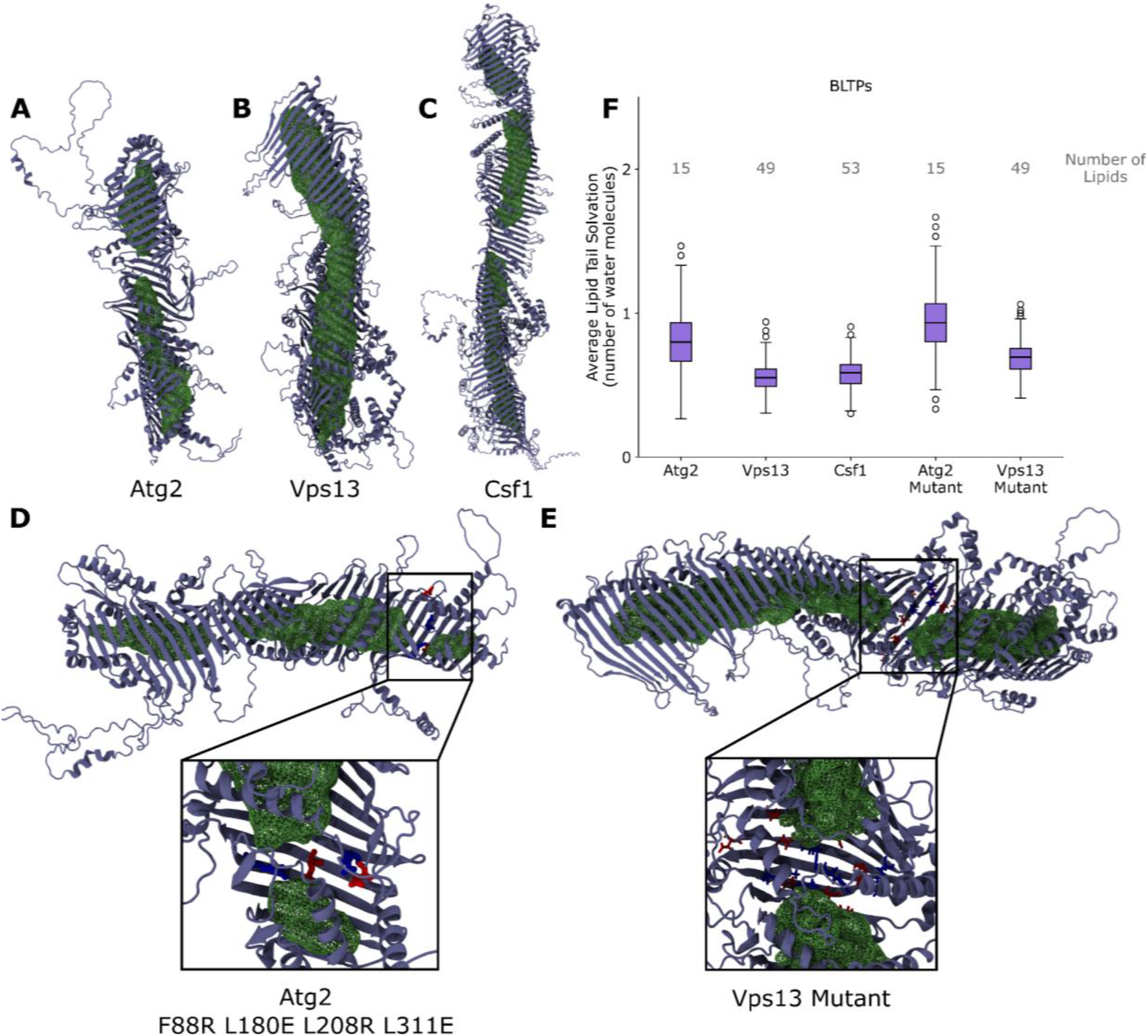
BLTPs possess a long hydrophobic cavity that can be filled with multiple lipids. **(A-E)** Spatial density maps and **(F)** lipid tail solvation (per lipid) using the iterative lipid addition process for BLTPs: **(A)** Atg2, **(B)** Vps13, **(C)** Csf1, and mutants of **(D)** Atg2 and **(E)** Vps13. Proteins are shown in purple; averaged lipid tail densities in green. The mutated residues are displayed in blue (arginine, lysine) or red (glutamate, aspartate).

Similar to the results described above, using an iterative lipid addition process we observed that the proteins can accommodate multiple lipids at the same time. Precisely, we could fit 15 lipids in Atg2, 53 in Csf1, and 49 in Vps13 with low tail solvation. Despite the large number of lipids in the cavity, the use of an elastic network to restrain the secondary structure prevents large protein conformational changes, thus indicating that the presence of multiple lipids is compatible with the initial AF models. A similar number of lipids (15) has been proposed for ATG2A, a human ortholog of Atg2 with the same length, using structural analysis^52^. The average solvation data of the lipid tails (Fig. 6F) are consistent with our previous analysis on shuttle LTPs (average < 1). Interestingly, even though the cavities are almost completely filled with lipids, the headgroups are arranged in such way that they face the solvent (Fig. S5). Further, the spatial density maps indicate a long hydrophobic conduit, but they also suggest the possible presence of bottlenecks in Atg2 and Csf1, which could be related to regulatory mechanisms.

Finally, to highlight the practical applications of our protocol, we designed mutations that block the hydrophobic cavity of Atg2, similar to those tested in ATG2A experiments^46, 53^. In these experiments, a small ring of hydrophobic residues was mutated to charged ones, resulting in impaired lipid transport *in vitro*. To mimic this approach, we selected four hydrophobic residues (F88, L180, L208, L311) close to the chorein motif of Atg2, where the lipid spatial density is large and uniform (Fig. 6D), and we mutated them to charged residues (arginine and glutamate). We next performed a 500 ns-long unbiased CG-MD simulation with the 15 lipids already inside the Atg2 mutant and computed their spatial density map at the end of the simulation. The resulting map (Fig. 6D) demonstrates that the new ring of charged residues generates a bottleneck where the lipids can no longer reside. A similar strategy has also been experimentally tested on Vps13^54^, showing that mutations in the middle of the hydrophobic cavity (V690D/L692R/L694E/I715K/A717D/M720K/I722D/I761R/I768E/F790D/M796D/L798R/ V802E/I816R/G820D/L827E) do not impair the lipid binding ability of Vps13, but do abrogate the Vps13 function in sporulation. As for the previous case, our protocol clearly indicates the formation of a significant bottleneck in the Vps13 mutant that could be responsible for the observed loss-of-function (Fig. 6E).

## Discussion

In the last few years, lipid transport by proteins has emerged as a central process in membrane and organelle homeostasis, but its molecular mechanisms remain largely unclear. To bridge this gap, we present here a computational protocol based on brute-force unbiased CG-MD simulations to propose a structural model for the binding pose of lipids inside LTPs. Our approach provides a physics-based hypothesis on the structure of the lipid-LTP complex that could contribute towards a mechanistic interpretation of *in vitro* and cellular experiments.

Our protocol is easy to reproduce (requiring only publicly available open-source software) and computationally cheap, as most simulations take only few hours to perform in a standalone GPU-accelerated workstation. In addition, it does not require previous knowledge of the binding pocket, and, by taking advantage of physicochemical properties of both lipids and proteins, can easily distinguish between polar and hydrophobic interactions, thus fully considering the amphipathic nature of most lipids. In addition, our iterative approach allows to propose a binding mode for LTPs binding multiple lipids in a dynamic way, with each lipid able to rearrange its localization inside the binding pocket upon the binding of subsequent lipids. This provides a clear advantage over docking methods, where the binding pose of each individual lipid is mostly static and can’t be easily modified upon the docking of subsequent molecules.

On the other hand, we acknowledge that our protocol has two main limitations. First, it is unable to discriminate between different lipids of similar sterical hindrance. As such, it can’t be used to investigate lipid specificity of LTPs. We foresee that future improvements in CG force fields or the use of multiscale strategies (e.g. by backmapping to all-atom simulations), possibly coupled with free-energy calculations, could help in this direction. Second, our protocol displays false negative results, as we could not straightforwardly identify any lipid binding pose for a few well-characterized LTPs, rather requiring *a posteriori* improvement to the protocol. This is very likely a consequence of the conformational plasticity of LTPs, that can adopt different conformations in their *apo* and *holo* states^55^. Hence, if the initial protein structure is in a “closed” conformation (for example in the presence of a lid that precludes lipid entry), our protocol is unable to identify the correct binding pathway and pose, since our CG approach can’t reproduce protein conformational flexibility as the protein structure is restricted by an elastic network. We expect that enhancing conformational sampling to generate a diverse set of starting protein structures, e.g. by all-atom MD simulations^55^ or by machine-learning based approach^56, 57^ could help mitigate this issue in the future.

In contrast, a strong advantage of our method is its robustness towards false positives. This suggests that whenever a de-solvated lipid binding pose is found, there is a high likelihood that this interaction has meaningful consequences for protein function, and that this warrants further experimental investigations. Specifically, we expect that our method will be extremely useful in at least two main areas. First, it will allow to generate mechanistic hypotheses for subsequent experimental validation for what concerns the molecular details of lipid transport, such as the identification of regulatory mechanisms including post-translational modifications or potential bottlenecks along the transport pathway. Second, we expect it will become a powerful tool for the direct identification of novel LTPs *in silico*, with the quality of the AI-based structural predictions as the main limiting factor.

## Methods

### Software details

MD simulations of all systems were performed with the GROMACS (v 2021.x)^58^ package, using the Martini 3 force field^59^. Molecular images were rendered using Visual Molecular Dynamics (VMD)^60^.

### System setup

The atomistic structures of the protein were obtained from the RCSB Protein Data Bank (PDB)^61^ and were converted to CG models using the martinize^62^ script. An additional elastic network with a force constant of 1000 kJ mol^-1^ nm^-2^ was used to restrain the secondary structure of the protein. A force constant of 300 kJ mol^-1^ nm^-2^ was also tested for Stard11 and FABP1. The proteins were then placed in a cubic box (in which the distance between the protein and the edge of the box was at least 2.0 nm), and one lipid molecule randomly placed in the bulk solvent. The system was then solvated and ionized with 0.12 M NaCl. For BLTPs, the structure was predicted using AlphaFold version 2.0 and, in the case of Vps13 and Csf1, different fragments were predicted and then aligned using common residues to get the structure of the entire protein. For LTPs that could bind multiple lipids in their cavity, the lipid addition procedure was repeated iteratively, such that the structure with n lipids bound in it served as the starting structure for the addition of (n+1)th lipid. This protocol was concluded once no more lipid entry into the hydrophobic cavity was observed.

### Simulation details for protein in water / LTP entry setup

Five independent replicas of 1 μs each were simulated for each protein-lipid system, with the exception of FABP1, Miga2, Stard11, Osh6 and Osh6 Δ69, for which five independent replicas were simulated for 3 μs each. Initial equilibration was carried out by performing energy minimization using the steepest descent algorithm, followed by a short MD run of 125 ps. Production runs were performed at 310K using a velocity-rescale thermostat^63^, with separate temperature coupling for protein and non-protein particles. The md integrator was used for the production runs, with a time step of 25 fs. The Parrinello-Rahman barostat^64^ was used to maintain the pressure at 1 bar, along with an isotropic pressure coupling scheme. The Coulombic terms were calculated using reaction-field and a cut-off distance of 1.1 nm. A cutoff scheme was used for the VdW terms, with a cut-off distance of 1.1 nm and the Verlet cut-off scheme for the potential-shift^65^. The non-bonded interactions were calculated by generating a pair-list using the Verlet scheme with a buffer tolerance of 0.005. The system setup and simulation parameters are in line with the recently proposed protocol for studying protein-ligand binding with the Martini force field^31^.

The Osh6 Δ69 protein was modelled by removing residues 36 to 69 that correspond to the lid of the ORP domain. Disulfide bridges in mutants NF1-SecPH 1603C-W1641C, and PITP V73C-210C were added using the CHARMM-GUI PDB Reader and Manipulator^66^ web server before coarse-graining the structure using the martinize script. The BLTP mutants were modeled with Pymol v2.3.0^67^ and aligned to the wild-type BLTPs filled with lipids after coarse-graining with the martinize script.

### Analysis

The minimum distance between the protein and the lipid was computed using the gmx mindist tool. An in-house tcl script was used to calculate the solvation number of the lipid, counting the number of water molecules within 5.0 Å of the lipid tail beads for each frame over the last 100 ns of the trajectory, for each replica.

Spatial density maps for lipids were computed with the Volmap plugin of VMD, by averaging over the last 100 ns of the trajectory of each replica.

To compute the distance to the lipid tails of the crystallographic ligands, the BB beads of the CG-MD trajectories were aligned to the N atoms of the protein crystal structure using the fit option of gmx trjconv. Then, a tcl script was used to compute the distance between the center of mass of the hydrophobic beads of the lipid and the center of mass of the lipid hydrophobic tails in the crystal structure. A similar procedure was used to compute the distance to the head groups of the ligands.

The lipid entry pathway into the LTPs during the MD simulation was visualized for each replica independently, and the percentage of replicas associated with the dominant lipid entry pathway was calculated by dividing the number of replicas in which the lipid entered the LTP via the dominant pathway, over the total number of replicas in which lipids entered the protein cavity overall. For example, in the case of MlaC, lipid entry occurred into the protein in 9/10 replicas, of which the entry occurred via the dominant pathway only in 7 replicas, hence yielding a percentage of 7/9*100 = 78% for this pathway.

## Acknowledgements

This work was supported by the Swiss National Science Foundation (PP00P3_194807 and PP00P3_163966). This work was also supported by grants from the Swiss National Supercomputing Centre under project ID s1030, s1132 and s1221 and has also received funding from the European Research Council under the European Union’s Horizon 2020 research and innovation program (grant agreement no. 803952). DA acknowledges support from the Margarita Salas program 2021-2023 funded by Ministerio de Universidades (MU-21-UP2021-030-53773022)

## Author contributions

S.S., D.A.L and S.V. designed research; S.S. and D.A.L. performed the simulations and analysed the data under the supervision of S.V. The paper was written by all authors

