## Supplementary Information for "Unbiased MD simulations characterize lipid binding to lipid transfer proteins"

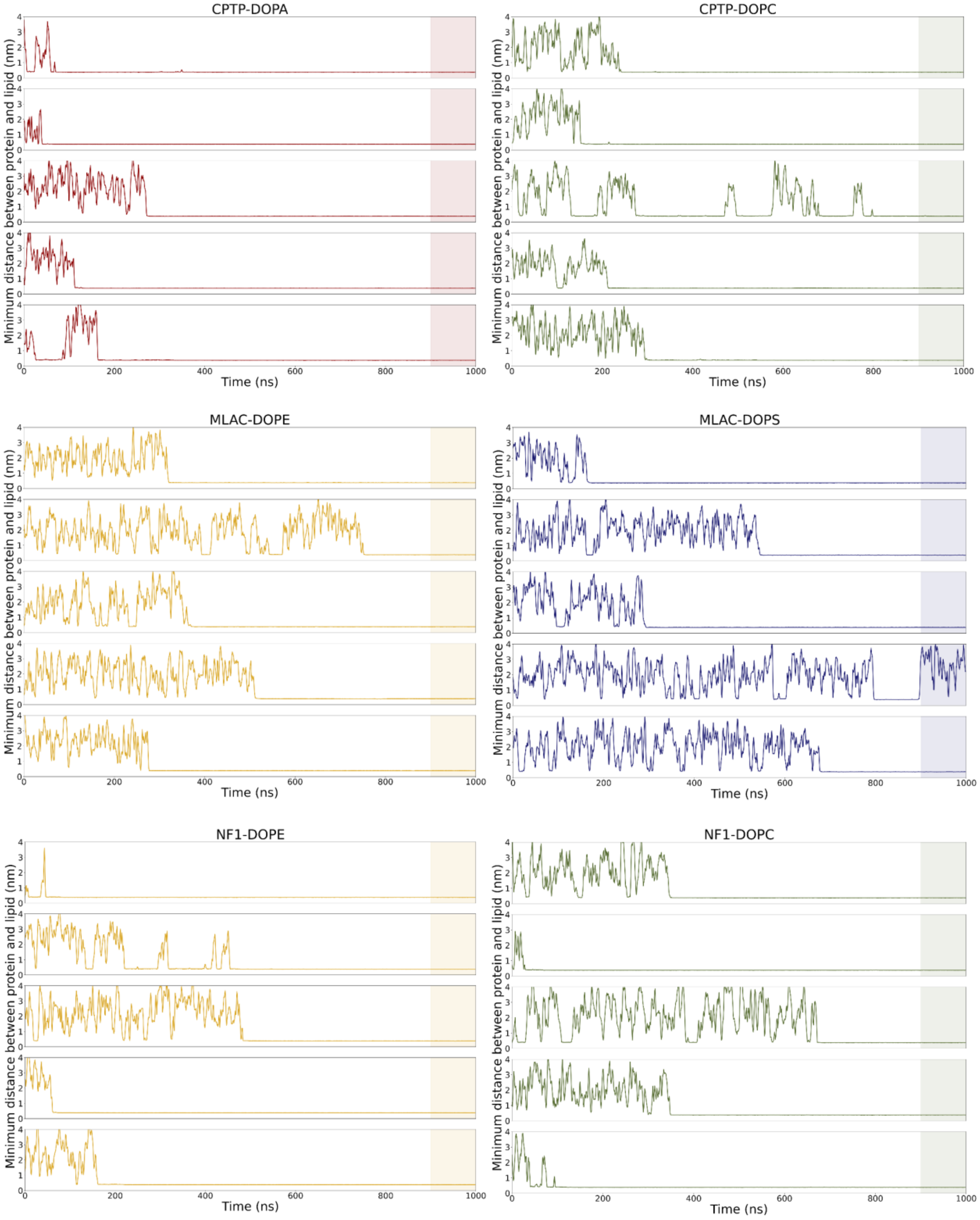

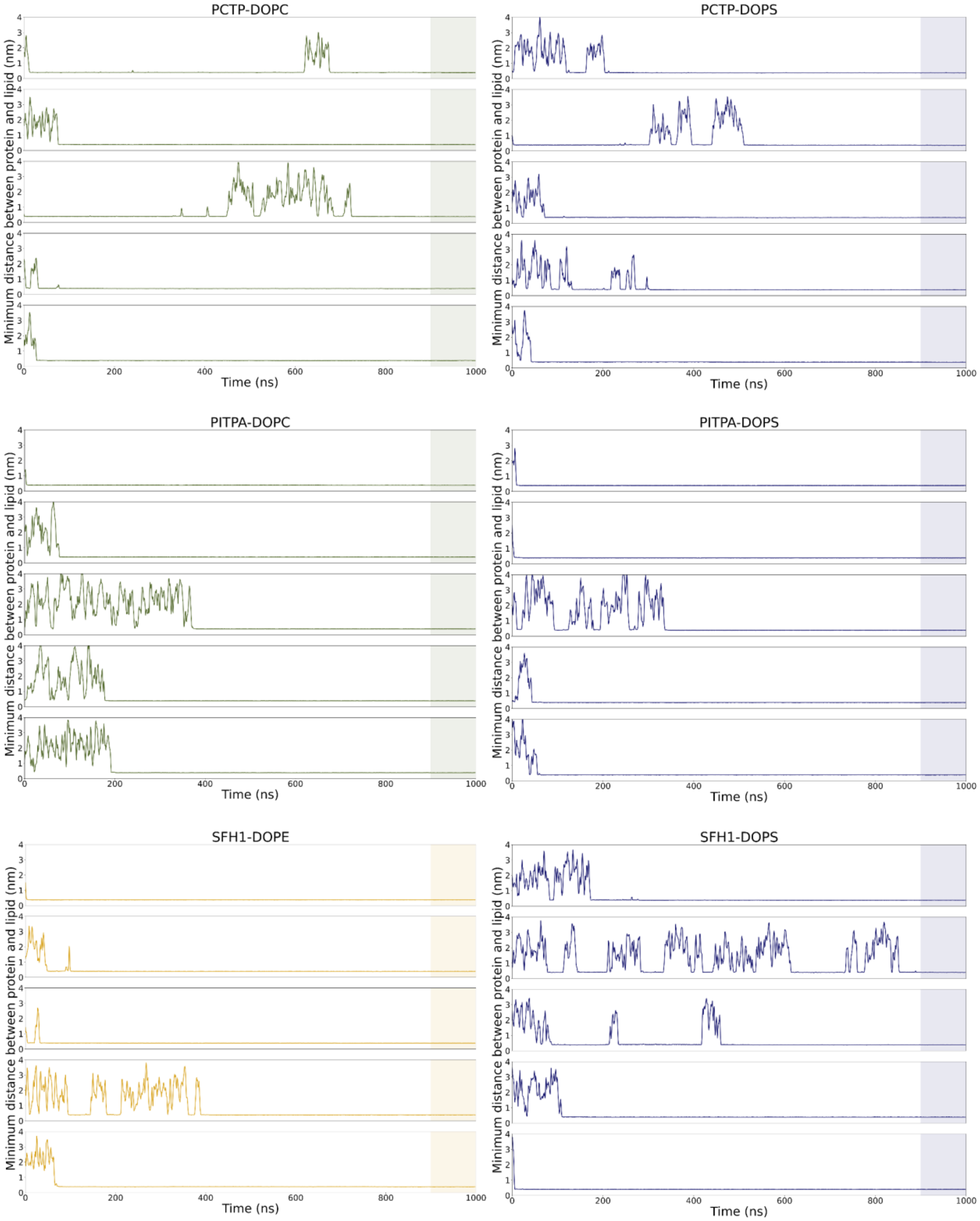
PCTP-DOPC
PCTP-DOPS
PITPA-DOPC
PITPA-DOPS
SFH1-DOPE
SFH1-DOPS
Minimum distance between protein and lipid (nm)
Time (ns)
0
200
400
600
800
1000
0
1
2
3
4

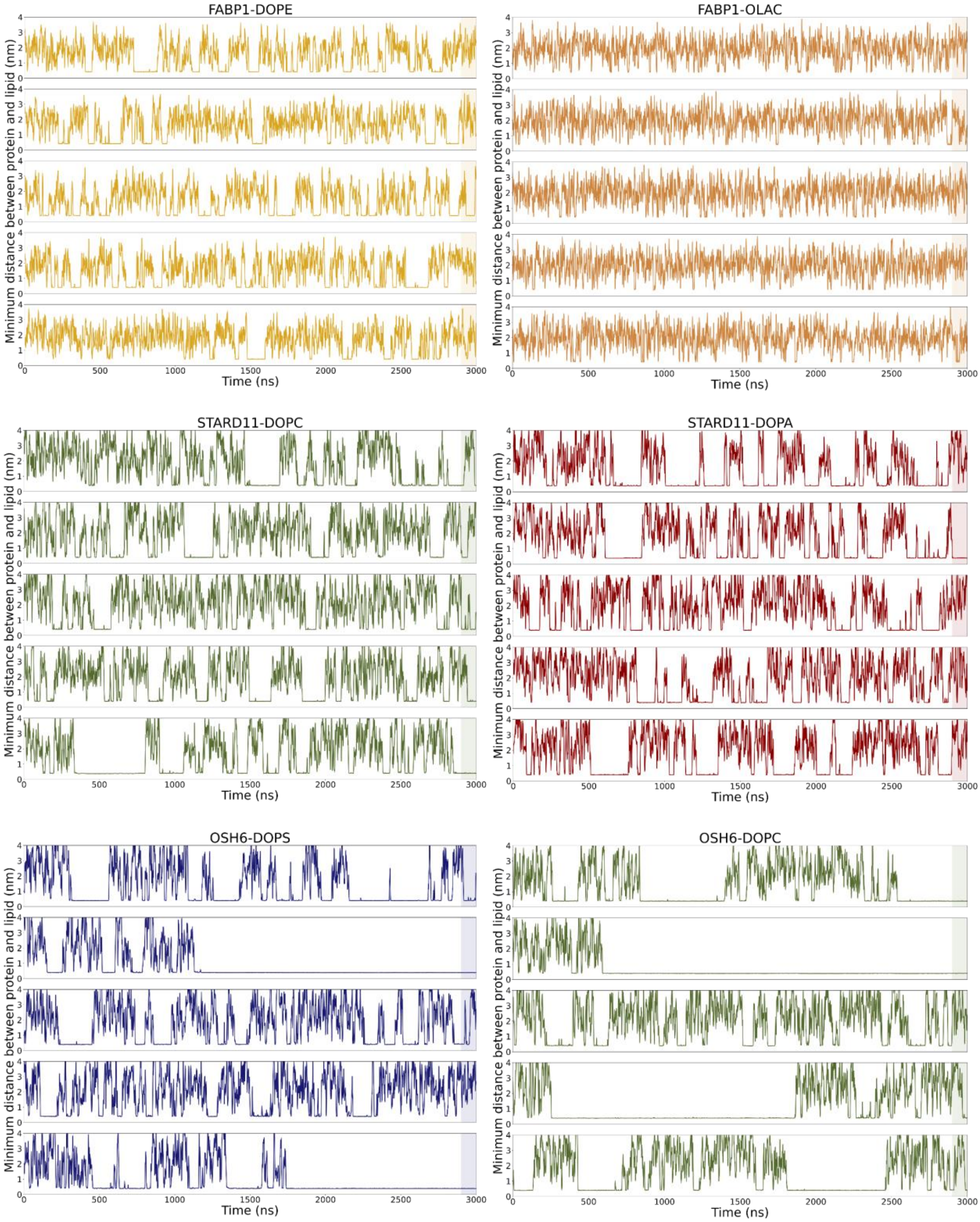
FABP1-DOPE
FABP1-OLAC
STARD11-DOPC
STARD11-DOPA
OSH6-DOPS
OSH6-DOPC
Minimum distance between protein and lipid (nm)
Time (ns)

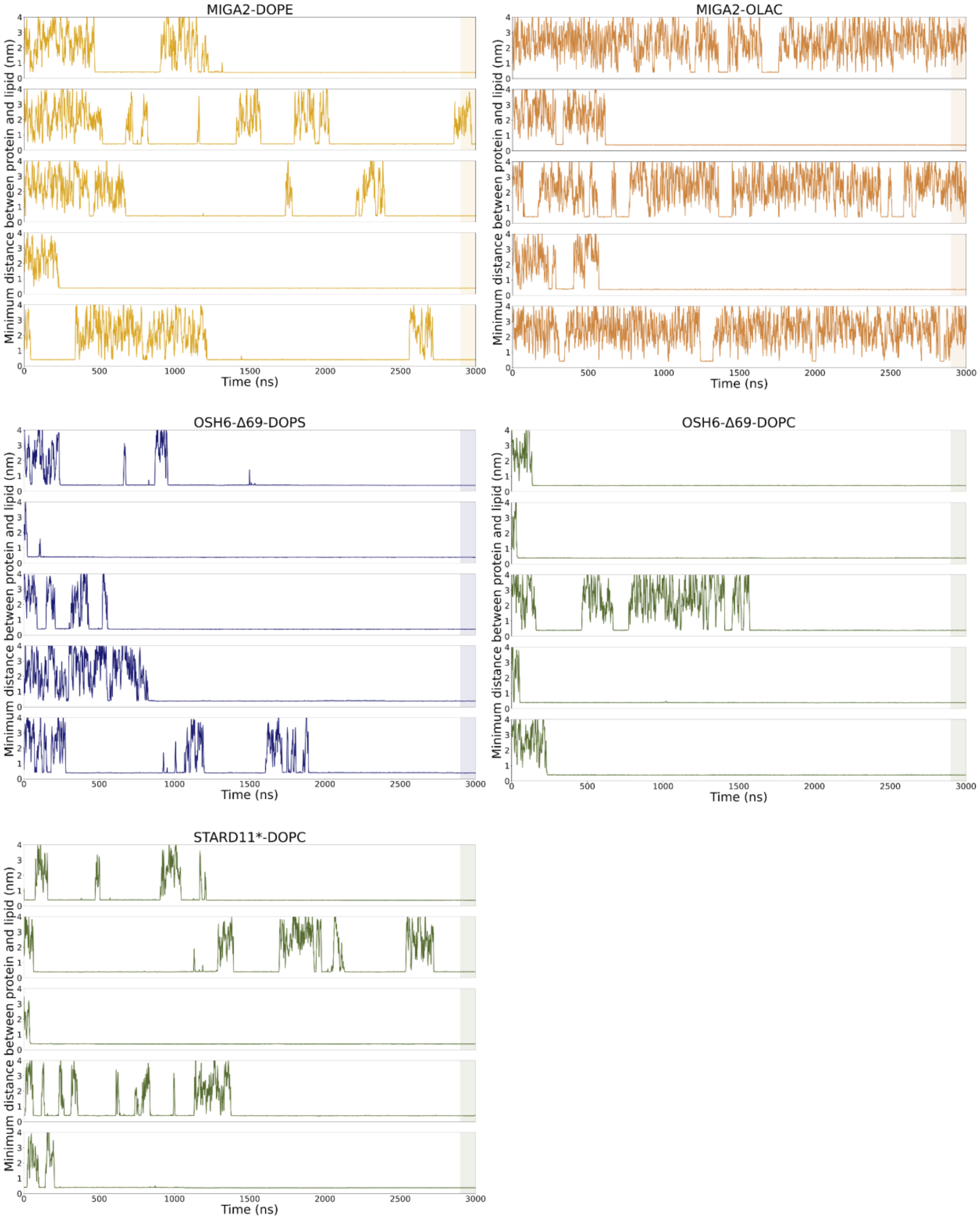


Figure S1: Time trace of minimum distance values between LTDs and lipids.

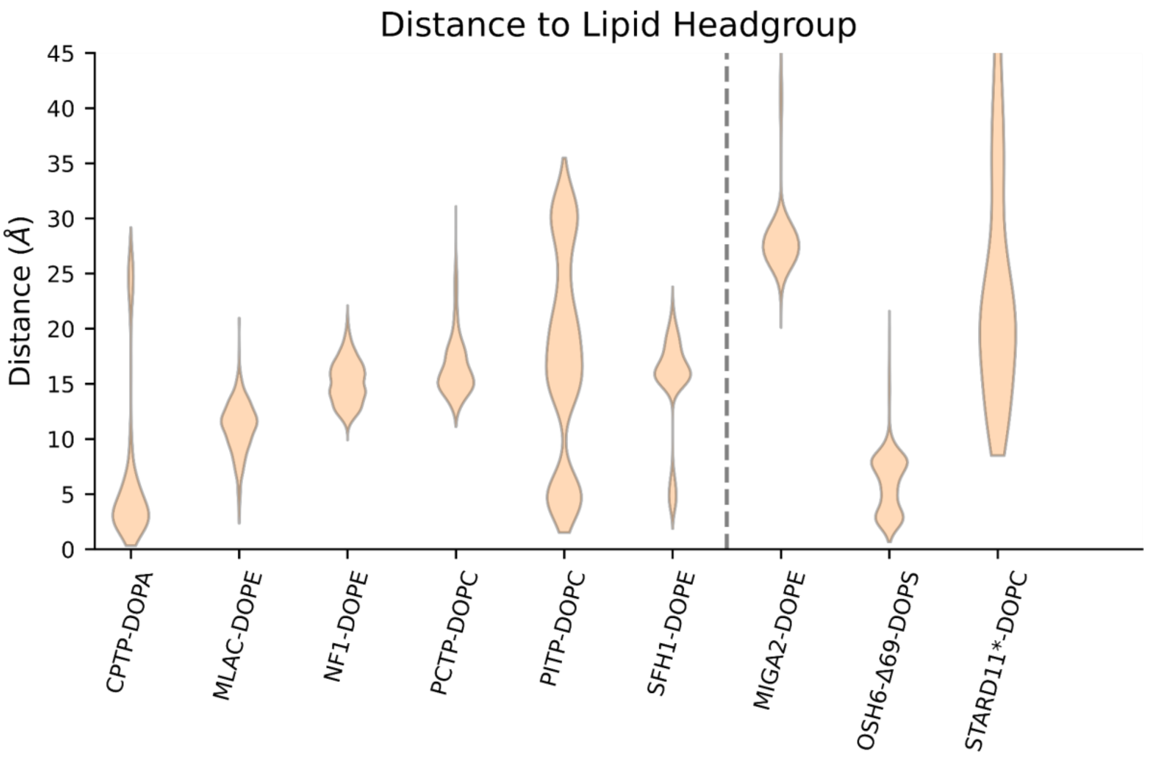


**Figure S2. Distance between the center of mass of the head group of the experimental ligands and the PO4 bead of the lipids tested in the simulations.** Only the frames with a lipid tail solvation of less than 2 were considered.

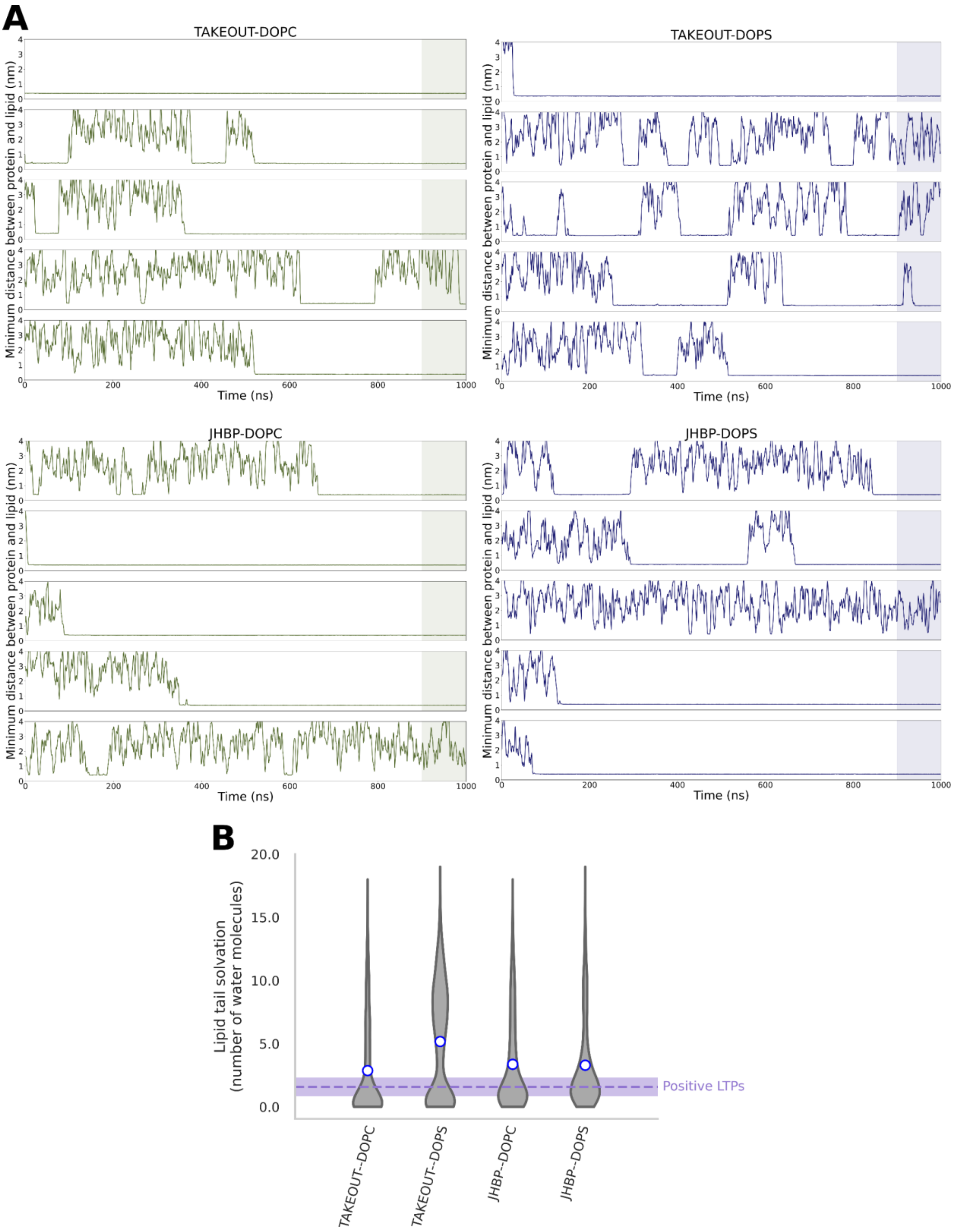


**Figure S3. Lipid binding to Takeout and Juvenile Hormone Binding Protein (JHBP) proteins. (A)** Time trace of minimum distance values between the lipids and the Takeout and JHBP proteins. **(B)** Lipid tail solvation in the last 100 ns of the trajectories.

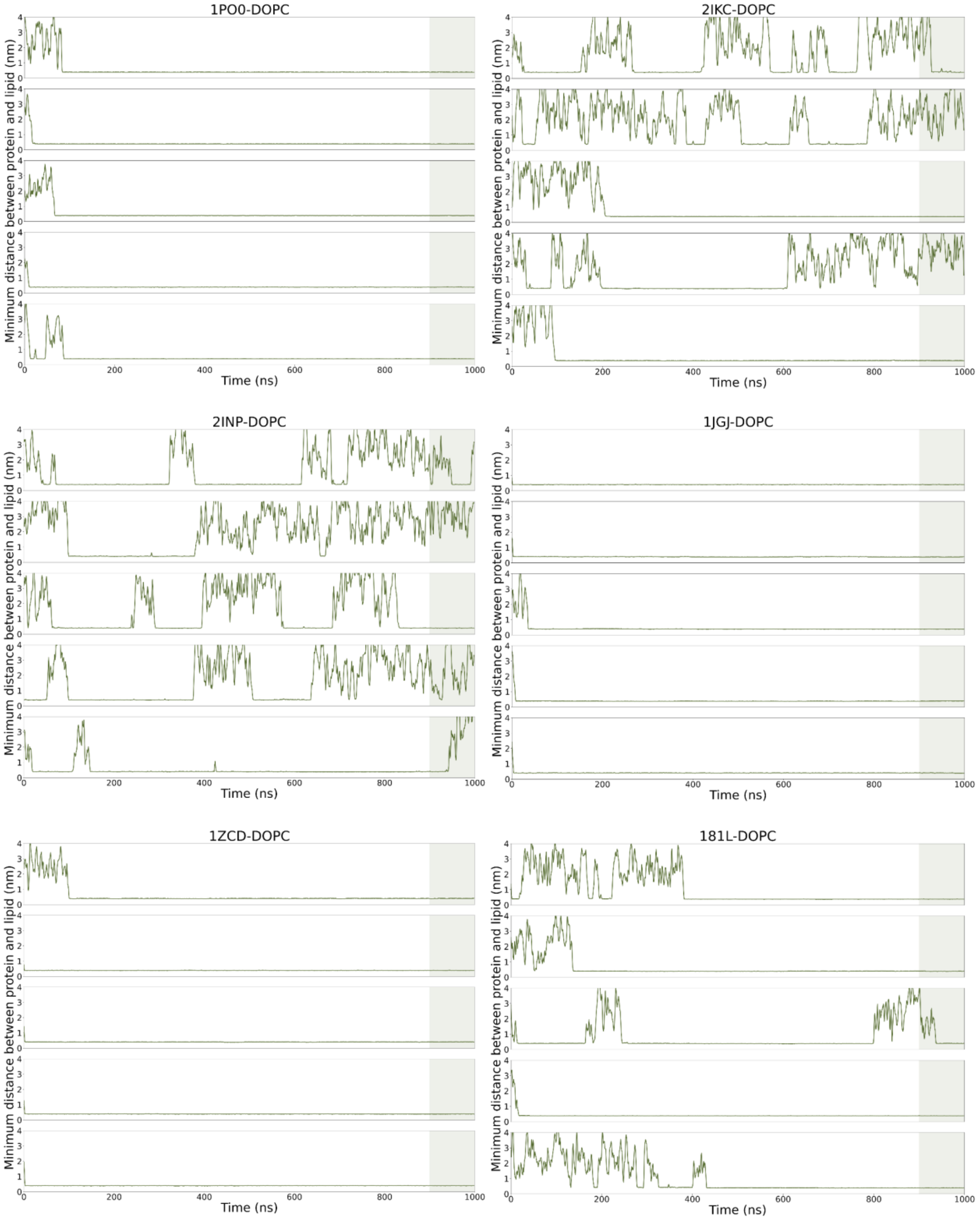
1PO0-DOPC
2IKC-DOPC
2INP-DOPC
1JGJ-DOPC
1ZCD-DOPC
181L-DOPC
Minimum distance between protein and lipid (nm)
Time (ns)
0
1
2
3
4
200
400
600
800
1000

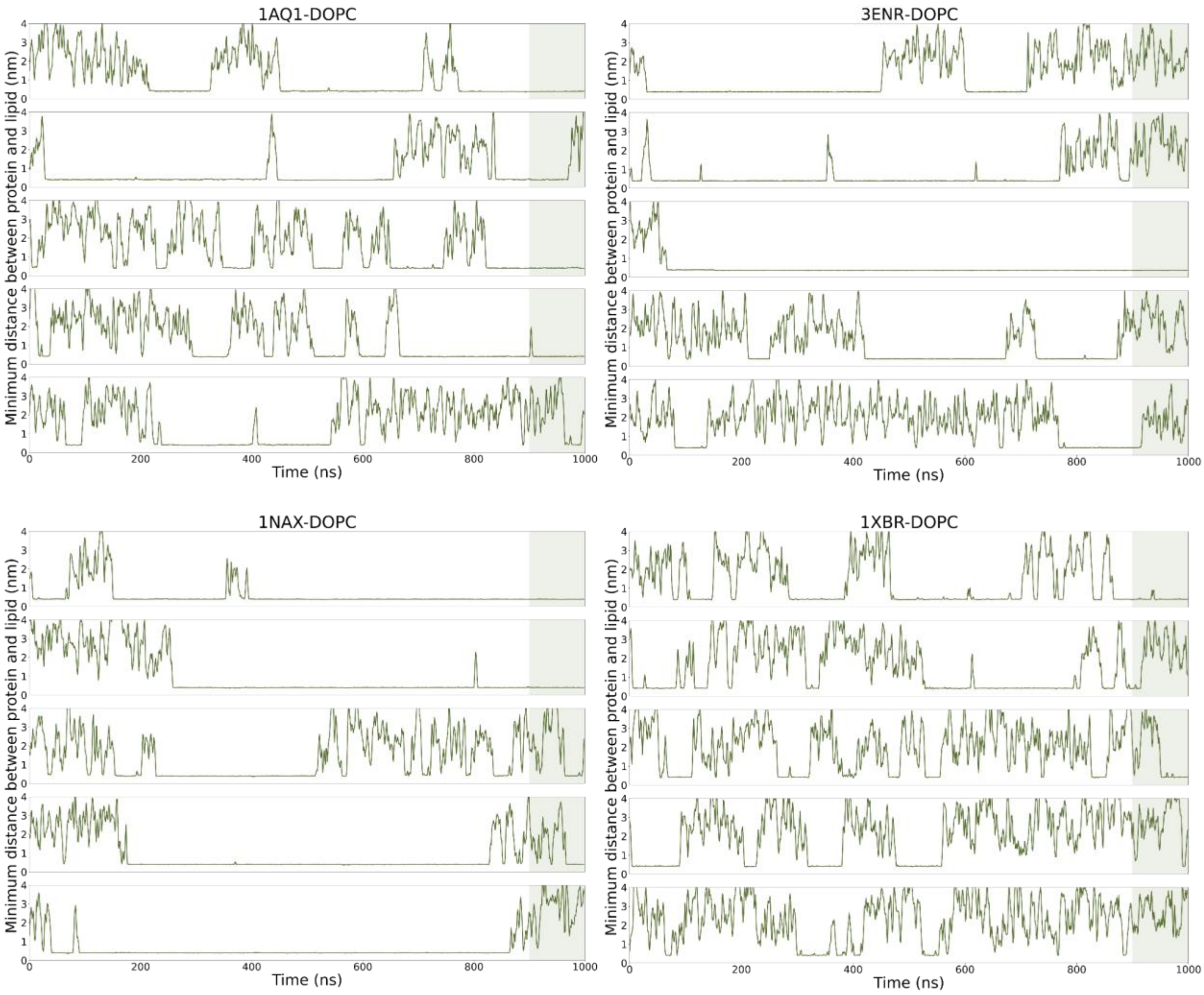


Figure S4: Time trace of minimum distance values between negative control proteins and DOPC.

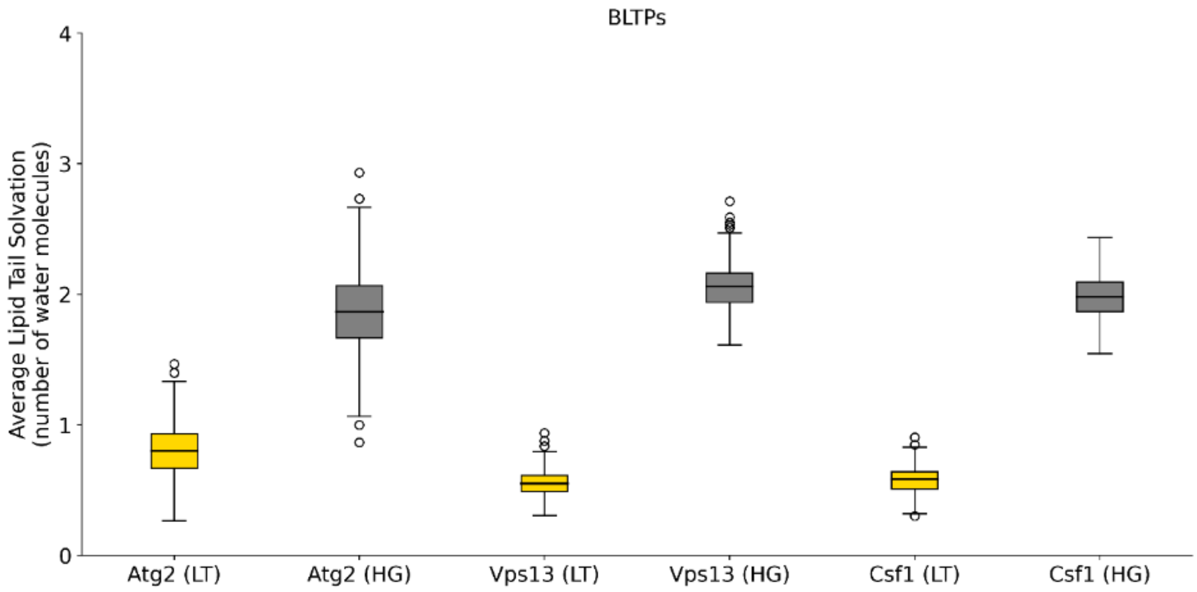


Figure S5. Comparison of the lipid solvation number of the lipid tails (LT) and the head groups (HG) for the BLTPs considered in this study (Atg2, Vps13, Csf1).
